# Landscape genomic prediction for restoration of a *Eucalyptus* foundation species under climate change

**DOI:** 10.1101/200352

**Authors:** Megan A. Supple, Jason G. Bragg, Linda M. Broadhurst, Adrienne B. Nicotra, Margaret Byrne, Rose L. Andrew, Abigail Widdup, Nicola C. Aitken, Justin O. Borevitz

## Abstract

As species face rapid environmental change, we can build resilient populations through restoration projects that incorporate predicted future climates into seed sourcing decisions. *Eucalyptus melliodora* is a foundation species of a critically endangered community in Australia that is a target for restoration. We examined patterns of genomic and phenotypic variation to make empirical based recommendations for seed sourcing. We examined isolation by distance and isolation by environment, determining gene flow up to 500 km and associations with environmental variables. Climate chamber studies revealed extensive phenotypic variation both within and among sampling sites, but no site-specific differentiation in phenotypic plasticity. Overall our results suggest that seed can be sourced broadly across the landscape, providing ample diversity for adaptation to environmental change. Application of our landscape genomic model to *E. melliodora* restoration projects can identify genomic variation suitable for predicted future climates, thereby increasing the long term probability of successful restoration.

## Introduction

Species around the globe face rapidly changing environments, often in combination with habitat degradation and fragmentation. These factors are expected to have a negative impact on biodiversity (Lindenmayer et al., 2010). Three processes enable species survival in the face of altered conditions-migration, adaptation, and phenotypic plasticity (Aitken & Whitlock, 2013; Aitken et al., 2008; Hoffmann et al., 2015; Nicotra et al., 2010). An important conservation strategy is to assist these natural processes to help build more resilient communities. We can help species shift to regions with their preferred environmental conditions by assisting migration of gene pools across the landscape (Aitken & Whitlock, 2013; Aitken et al., 2008). We can aid populations to survive in situ by ensuring that sufficient genomic variation exists for adaptation to changing environments (Hoffmann et al., 2015). We can enable individuals to respond to a greater range of environments by conserving existing phenotypic plasticity (Nicotra et al., 2010).

Seed sourcing during landscape restoration provides an ideal opportunity to apply scientific knowledge to enable these key processes and improve conservation outcomes (Broadhurst et al., 2008; Prober et al., 2015). For example, seed sources can be selected to restore historical patterns of gene flow across a fragmented landscape, incorporate high genomic diversity, and/or increase phenotypic plasticity. Seed sources can also be matched with current or projected future climates, enabling assisted migration to favorable environments (Aitken & Whitlock, 2013; Williams et al., 2014).

Historically, restoration often focused on geographically restricted local sources of seed under the premise that this would improve restoration outcomes by reducing the risk of maladaptation to local conditions and preventing outbreeding depression (Broadhurst et al., 2008). However, there are several potential drawbacks to this narrow local focus. In a fragmented system, narrow local seed sourcing reduces the number of potential source populations, thereby reducing the pool of available genetic material. This reduced gene pool may result in inbreeding depression in future generations, especially if combined with small population size (Broadhurst et al., 2008). Obtaining potential seed sources from a wider geographical area can increase genomic and phenotypic diversity, thereby increasing the ability of the species to survive in situ (Broadhurst et al., 2008). Additionally, the focus on maintaining local adaptation in situ assumes a static environment, not the rapidly changing environment that occurs today. As local conditions change, traits and genes that may have conferred an advantage in the past might not be suitable in the future environment. In recent year, climate adjusted provenancing has been proposed, which is a seed sourcing strategy that incorporates climate variability and focuses on sourcing seed that is predicted to be adapted to future climates (Byrne et al., 2013; Prober et al., 2015). This strategic assisted migration of variation across the landscape can aid in the establishment of populations that are more adaptable to future environments (Prober et al., 2015).

To determine the appropriate seed sourcing strategy and to identify optimal seed sources for a reforestation project, empirical knowledge of genomic variation for the target species can provide valuable information. The technology now exists to assess genomic variation in any target species, enabling determination of patterns of Isolation By Distance (IBD) and Isolation By Environment (IBE). IBD is the association between genomic distance and geographic distance resulting from patterns of dispersal. IBE is the association between genomic distance and environmental distance, while controlling for geographic distance (Wang & Bradburd, 2014). Landscape genomic models can be generated by fitting geographic and environmental variables to the observed genomic diversity. These predictive models can optimize the genetic material selected for restoration and should improve long term outcomes (Hoffmann et al., 2015; Williams et al., 2014).

The extent of phenotypic plasticity in potential seed sources can be measured in growth assays of seedling traits across contrasting environmental conditions. The magnitude of the environmental response can be compared among maternal lines or populations and may identify populations that differ in their response to the environment. Such differing responses have been seen in some species of *Eucalyptus* (Andrew et al., 2010; Byrne et al., 2013; McLean et al., 2014).

*Eucalyptus melliodora* (A.Cunn. ex Schauer), commonly called yellow box, is an iconic Australian species that is the subject of extensive restoration efforts across its distribution. It is a foundation species of a critically endangered ecological community: the White Box–Yellow Box–Blakely’s Red Gum Grassy Woodland and Derived Native Grassland (Department of Environment and Climate Change and Water, 2011; Department of the Environment and Heritage, 2006; Threatened Species Scientific Committee, 2006). This woodland community exists in a fragmented landscape, with less than 5% of its original distribution remaining, mostly in small remnant patches (Department of Environment and Climate Change and Water, 2011; Department of the Environment and Heritage, 2006; Threatened Species Scientific Committee, 2006). Efforts to restore this endangered woodland community are ongoing and restoration practitioners are seeking scientific recommendations to improve seed sourcing. Climate change is an important consideration in seed sourcing decisions because species distribution modelling predicts that most eucalypts will need to shift their distributions considerably in response (González-Orozco et al., 2016). In particular, for *E. melliodora* ecological niche modelling predicts that by 2090 the species distribution will shift toward the southeast and suitable areas will decrease by 77% as a result of environmental changes (Broadhurst et al., *in review*).

Here we survey genomic variation in 275 individuals from 37 sites across the present range of *E. melliodora*. We fit the genotypic data to geographic distance and key environmental variables at the site of origin. We find that effects of genomic isolation by distance begin at approximately 500km. This empirical estimate of "local" is much farther than what is often practiced for local provenancing. We also find that features of the abiotic environment can further explain genomic differentiation after accounting for geographic distance. We also examine seedling growth characteristics under simulated climate conditions and find significant variation in growth traits both within and among sites, but no significant variation in phenotypic plasticity across sites. Our landscape genomic model, which can empirically define local provenances and identify variation suitable for predicted future climates, can help build resilient populations through scientifically based restoration.

## Results

### Genotyping by Sequencing

We selected leaf material from 39 sites, sampling 3-10 trees per site (Supplemental Table S1). For each sample we Illumina sequenced a Genotyping by Sequencing (GBS) library (Elshire et al., 2011) and used a reference alignment-based approach to call genotypes. We conducted a preliminary analysis, based on 123,227 SNPs and removed 69 samples due to greater than 60% missing data. Visual examination of a cluster dendrogram of genomic distance between samples showed that technical replicates cluster closely together (Supplemental Fig. S1). A preliminary principal coordinate analysis (PCA) identified 19 samples that were strong genomic outliers (Supplemental Fig. S2), likely misidentified samples or recent hybrids. This result is consistent with minor morphological differences noted in these samples, as well as previous microsatellite work (Broadhurst et al., *in review*). After removal of poor quality and geographic and genomic outlier samples, we reran the genotyping with the remaining 280 samples, resulting in 9,781 SNPs after filtering. A second preliminary PCA identified an additional 5 outlier samples that we considered sufficiently differentiated from the main *E. melliodora* cluster to merit removal for downstream analyses (Supplemental Fig. S3). We removed these samples and reran the missing data filter. The final data set included 275 samples from 37 sites (Fig. 1A), genotyped at 9,378 physically distinct SNPs (>300 bp apart).

**Figure 1.**
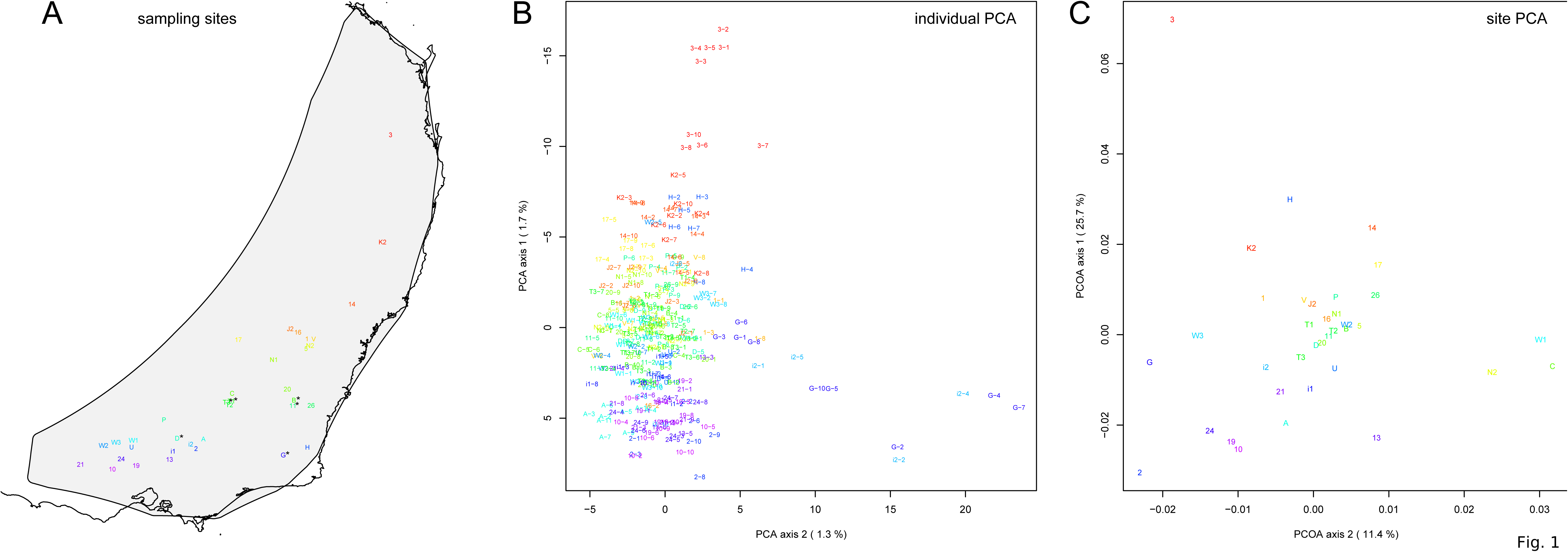
Map of sampling sites and PCA of genomic distance between samples. (A) A map of the geographic locations of the 37 sampling sites in southeastern Australia. Sampling locations are labeled with the site name and color coded in a rainbow gradient based on latitude. Black asterisks indicate 6 sites used for growth chamber experiments. The gray background shading indicates the species distribution polygon. (B) Principal coordinate analysis of the genomic distance between individual samples. Samples are labeled with a sample name that indicates the site name and tree number. Samples are color coded by site to match the map. The percentage on each axis indicates how much of the genomic variation between individuals was explained by the axis. (C) Principal coordinate analysis of F_st_ between sampling sites. Sampling sites are labeled by name and color coded to match the map. The percentage on each axis indicates how much of the variation in F_st_ between sampling sites was explained by the axis.

### Genomic Analyses

A PCA of genomic distance among samples showed continuous variation with little suggestion of discrete population structure (Fig. 1B). The samples largely formed a single cluster, with the first PCA axis corresponding roughly to latitude. Outside of the main cluster, samples from the northernmost site separated out along the first PCA axis (y-axis) and a few samples from two other sites separated out along the second PCA axis (x-axis). Together, the first two PCA axes explained 3.0% of the genomic variation among individuals. The Mantel test estimated that the natural log of the geographic distance between samples explained 2.3% of the variation in individual genomic distance, indicating weak, but statistically significant, isolation by distance (p=0.0001). We summarized genomic diversity between sampling sites using pairwise F_st_. For all comparisons F_st_ was low (mean F_st_=0.04, sd=0.02) (Supplemental Table S2). The maximum F_st_=0.10 occurs between sites 3 and 13, which are separated by over 1200 km. Similar to the PCA of genomic distance among samples, the PCA of F_st_ between sampling sites corresponded roughly to latitude (Fig. 1C). In contrast, the first two axes of the PCA of F_st_ between sampling sites explained a higher percentage of variation (37.1%). These results highlight the tremendous amount of genomic variation within sampling sites, as well as the ability of thousands of independent genomic markers to distinguish between more distant sampling sites.

The site by site pairwise F_st_ matrix was used to test for geographic and environmental associations using generalized dissimilarity modelling (GDM) (Ferrier et al., 2007; Fitzpatrick & Keller, 2015; Thomassen et al., 2011). Of the 28 environmental variables considered for the model, we removed 12 variables because the single variable model explained less than 5% of the deviance (bioclimatic variables 2, 5, 6, 9, 10, 14, 17, 19; elevation; water at depth; Prescott Index; and MrVBF). We removed an additional 9 variables due to high correlation and lower explanatory power than another remaining variable (bioclimatic variables 1, 4, 7, 12, 13, 15, 18; surface nitrogen; and surface phosphorus) (Supplemental Table S3). We ran permutation testing on a model with the remaining 7 variables and geography. This highlighted an additional 2 variables with low statistical significance and low explanatory power that we removed from the final model (surface water and bioclimatic variable 8). We also removed phosphorus at depth because, although it explained a substantial amount of genomic variation, the sampled sites were not well distributed across the range of values.

As a result, we included four environmental variables in the final model: isothermality (bioclim 3), mean temperature of the coldest quarter (bioclim 11), precipitation of the wettest quarter (bioclim 16), and total soil nitrogen at 100-200 cm (nitrogen at depth) (Supplemental Fig. S4). The GDM model with these four variables plus geographic distance explained 40% of the variation in sampling site genomic differentiation (F_st_). The GDM model showed a positive non-linear relationship between environmental distance and genomic distance (Supplemental Fig. S4A). To test the predictive power of the GDM model, we used a cross validation approach by generating 1000 models with a random 30% of sampling sites removed. GDM proved satisfactory at predicting genomic differences between removed sites (cross validation correlation mean=0.73, standard deviation=0.12) (Supplemental Fig. S4B).

Of the four environmental variables, nitrogen at depth showed the strongest relationship with genomic distance, with changes in genomic distance predicted across the range of nitrogen values (Supplemental Fig. S4C). Mean temperature of the coldest quarter was the second strongest predictor, showing a similar pattern as nitrogen (Supplemental Fig. S4D). Precipitation of the wettest quarter was the third strongest environmental predictor, predicting the largest change in genomic distance between 250 and 400 mm (Supplemental Fig. S4E). Isothermality (mean diurnal range divided by annual temperature range) was the final predictor, predicting the most change in genomic distance at higher values (Supplemental Fig. S4F).

Geographic distance showed a non-linear relationship with genomic distance. The geographic spline predicted no genomic differentiation until close to 500 km, at which point an increase in geographic distance predicted an increase in genomic distance (Fig. 2). Randomly subsampling sites showed that the predicted genomic distance for large geographic distances was quite variable, but for sites less than 500 km apart, all iterations consistently predicted little genomic differentiation between sites (Supplemental Fig. S4H).

**Figure 2.**
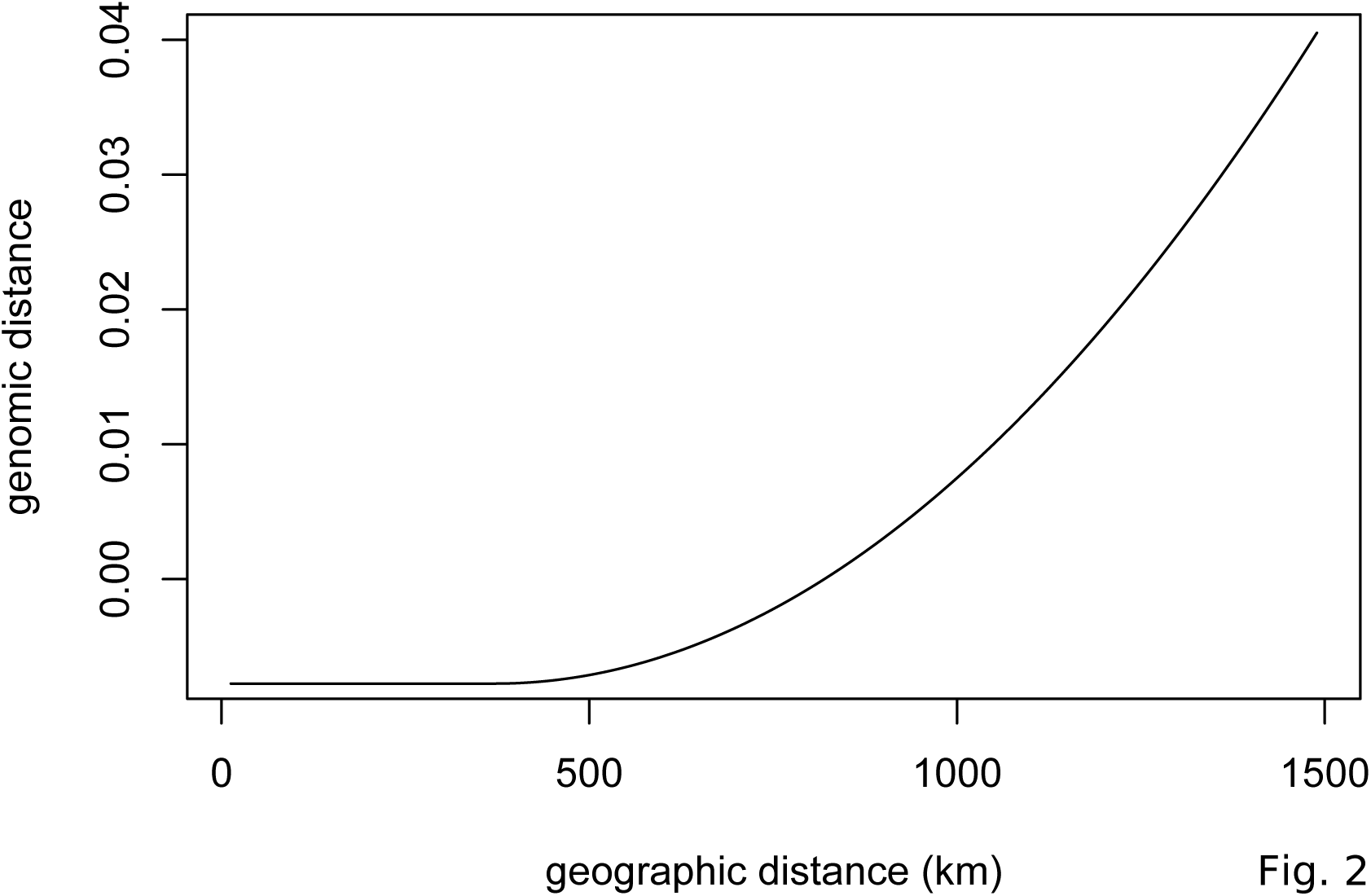
Estimated genomic variation as a function of geographic distance. The geographic spline estimated from the GDM model showing little predicted genomic change between sites less than 500 km apart and increasing genomic variation as geographic distance increases beyond 500 km.

To project the final GDM model onto the current environmental landscape, we first delineated the geographic extent of the analysis by defining an *E. melliodora* distribution polygon. We then projected the GDM model onto this region using the current values of the environmental variables across the landscape. This analysis partitioned the landscape into a number of regions with different predicted genomic compositions, including northern coastal, northern inland, and southern regions (Fig.3A). While the biggest differences occurred in regions with few sampling sites, there is a distinction between the northern and southern sites, as well as between sites on opposite sides of the Great Dividing Range in the southern region (e.g. site G versus site A, Fig. 3A). These projections highlight where environmental filtering of genotypes may have occurred due to different selective pressures.

**Figure 3.**
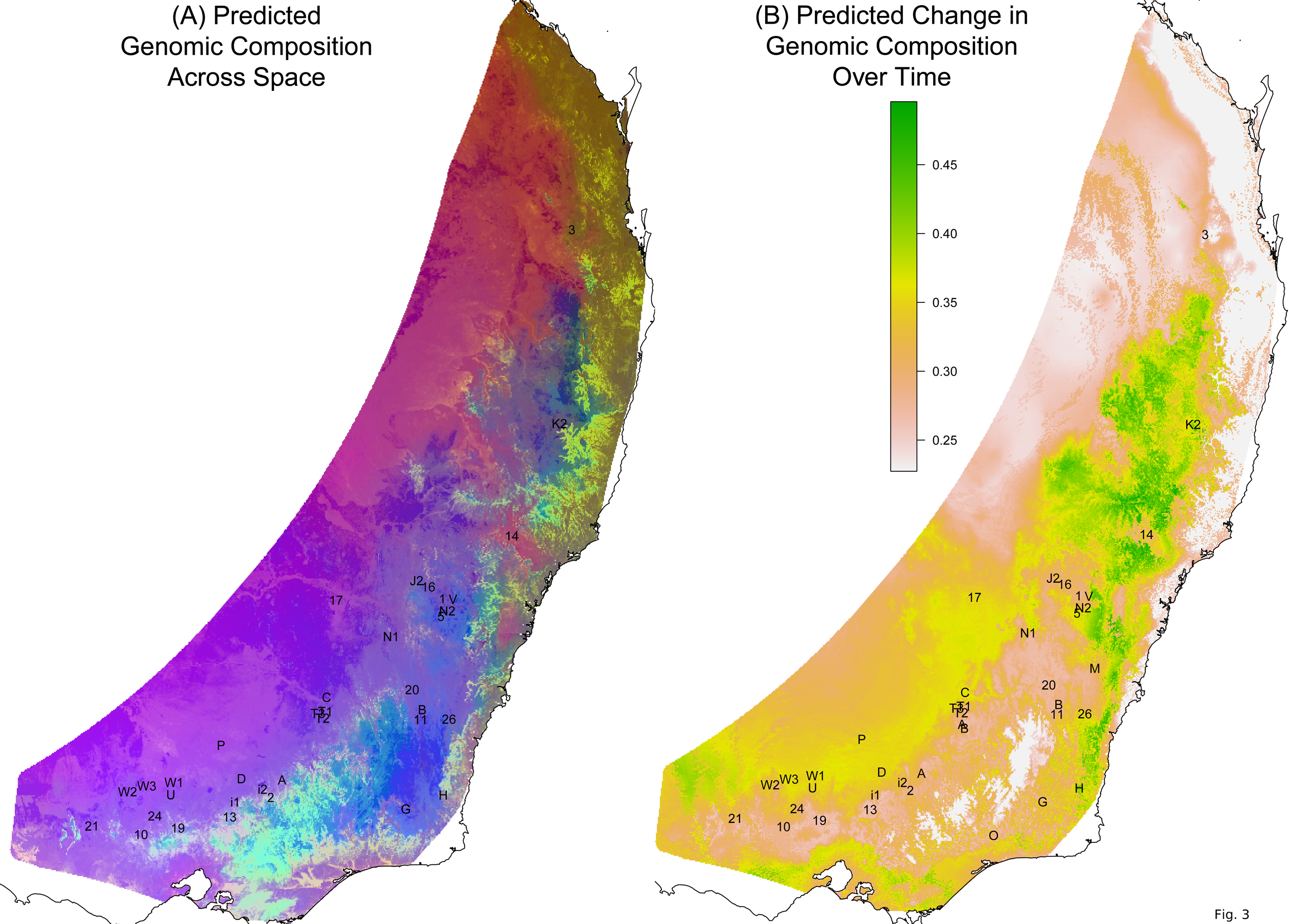
Predicted spatial and temporal variation in genomic composition. (A) The spatial distribution of predicted genomic variation based on projecting the GDM model onto geography and current environmental conditions. Regions with similar colors are predicted to have similar genomic composition. (B) The predicted temporal genomic variation based on comparing the GDM model projected onto current environmental conditions and predicted environmental conditions for 2070. The higher the difference (green colors), the more genomic change predicted between current and 2070 conditions. Sampling sites are labeled in black text.

We compared the GDM model projected onto current conditions to the GDM model projected onto 2070 climate predictions. This analysis scored each position across the landscape based on how much genomic change was predicted to occur in response to changing environmental conditions (Fig. 3B). For the middle north region (around sites K2 and 14) and the southern areas towards the coast, the models predicted more intense natural selection in response to climate change. Thus, these areas could be prioritized for assisted migration.

We also used the GDM model to compare the genomic composition under future environmental conditions at a single location to the genomic composition under current climate conditions across the landscape. This comparison is useful for identifying optimal seed sources for restoration sites given climate change scenarios. We demonstrated this utility by selecting two hypothetical reforestation sites and identifying distinct regions that would provide favorable seed sources for each site (Fig. 4). The analysis for the southern reforestation site identified a large portion of the southern distribution, centered at the reforestation site. For this site it appears that the selected areas are largely driven by the pattern of isolation by distance, in particular the lack of genetic differentiation for long geographical distances. The analysis for the northern reforestation site identified a more limited range of areas across the landscape, possibly driven in part by a decreased power due to lower sampling intensity in the north. In addition to identifying a narrow region in the north that is centered on the reforestation site, a number of more distant areas along the coast were also identified, indicating these selected areas are driven more by patterns of isolation by environment than isolation by distance.

**Figure 4.**
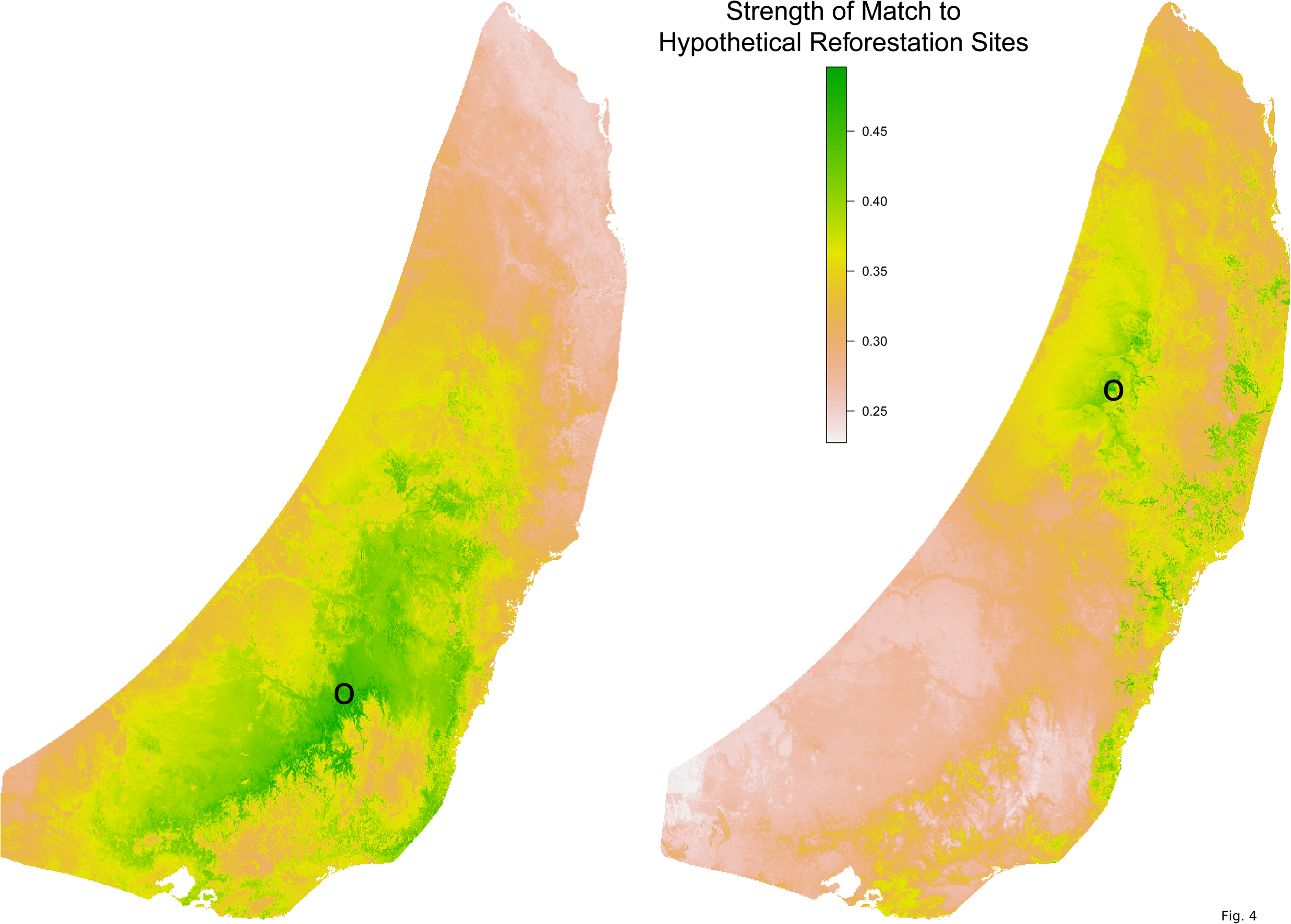
Optimal seed sourcing locations for hypothetical reforestation sites. The predicted genomic similarity of hypothetical reforestation sites (indicated by black circles) to potential seed sourcing locations under a climate change scenario for 2070. Dark green areas indicate seed sourcing areas predicted to best match future conditions at the hypothetical reforestation sites; white and brown areas indicate areas of potential genomic mismatch.

These genomic analyses suggest that for woodland restoration a geographically wider and environmental model based approach to seed sourcing would allow incorporation of more genetic diversity and enable better matching of the selected genotypes to current and predicted future environmental conditions at the reforestation site.

### Growth Experiments

We conducted a climate controlled growth experiment to measure variation in seedling growth traits among sampling sites and assay phenotypic plasticity. We grew seedlings from six sites, with six maternal lines per site, at two different climate regimes (average summer conditions and 5°C hotter summer conditions). For analysis of seedling height and total leaf length, we analyzed a total of 291 seedlings (from 32 maternal lines representing six sampling sites) that were determined to be well established at the five week measurement. For analysis of the relative height increment, we analyzed a total of 560 seedlings (from all 36 maternal lines) for which were able to calculate this metric. There were four seedlings that were outliers for the relative growth increment (>0.035). These outliers had little effect on the results of the linear models, so we included them in the final analysis.

The models for all three response variables (seedling height, total leaf length, and relative height increment) showed that all fixed effects (sampling site, maternal line nested within sampling site, and experimental condition) were statistically significant at the p=0.05 level (Supplemental Table S4). Experimental condition explained a small percentage of the variation (1.2-8.1%), as did sampling site (1.8-17.7%) (Supplemental Fig. S5). Maternal line tended to explain a larger amount of variation (10.6-27.6%).

However, most of the variation remained unexplained (56.6-71.5%). None of the three response variables showed significant variation in phenotypic plasticity across sites (all maternal line/sampling site by experimental condition interactions p>0.50) (Supplemental Fig. S6 and Table S5).

We then conducted an outdoor drought experiment using a subset of seedlings from the temperature experiment. We analyzed 146 seedlings representing 20 maternal lines from five sampling sites. These seedlings were grouped into 73 pairs, with one of each pair assigned to each treatment—well watered versus drought. We analyzed variation in four response variables: stomatal conductance, leaf length to width ratio, relative chlorophyll content (SPAD index), and specific leaf area (SLA, leaf area divided by dry mass).

The droughted seedlings had significantly lower stomatal conductance rates than the well watered ones, indicating that the seedlings were stressed (p<0.00001) (Supplemental Table S6). Treatment explained most of the variation in stomatal conductance (62.3%), while maternal line and sampling site explained only a small amount of variation (5.8% and 0.9% respectively) (Supplemental Fig. S7). For the remaining three response variables (leaf length to width ratio, SPAD, and SLA), much of the variation was unexplained (40.5%-70%). Experimental condition was not statistically significant and explained little to no variation (0.0-4.4%) (Supplemental Fig. S7 and Table S6). Sampling site and maternal line were statistically significant in the linear models at the p=0.05 level and explained some variation (6.7-21.2%) (Supplemental Fig. S7 and Table S6). Smaller, thicker leaves, and thus lower SLA values, were expected for droughted seedlings and for seedlings grown from seed collected from drier areas. Consistent with this expectation, the seedlings subjected to drought conditions showed lower SLA values. However, seedlings from drier sampling sites (D and T3) showed higher SLA values than more mesic sites (B, G, and 11), contrary to expectation (Supplemental Fig. S7). None of the four response variables showed significant variation in phenotypic plasticity across sites (all maternal line/sampling site by experimental condition interactions p>0.13) (Supplemental Fig. S8 and Table S7).

In addition to measuring seedling growth traits, we also examined the shape of the leaves from the drought experiment seedlings. We noted substantial variation in leaf shape, both among sites and within (Supplemental Fig. S9). The remarkable amount of phenotypic variation in the seedlings is consistent with the high levels of genomic variation measured both among sites and within sites.

## Discussion

*Eucalyptus melliodora* is a foundation species in a critically endangered woodland community that now occupies a fraction of its former distribution and is the subject of restoration projects across its native range. Our examination of the distribution of genomic and phenotypic variation across the range of this species provides valuable information for sourcing seed for restoration, including empirically defining local provenances and matching genotypes to predicted future environmental conditions.

Examining the relationship between genomic and geographic distance, we are able to empirically define "local" in this species to be on the order of 500 km, which is substantially farther than the current practice. This new definition encourages restoration projects to source seed more broadly across the landscape. In a highly fragmented landscape this will increase the number of potential source sites, potentially enabling the collection of higher quality seed with increased genetic diversity (Broadhurst et al., 2008). Incorporating more naturally occurring genomic variation can increase the adaptive potential of the restored population by providing the substrate for adaptation to rapidly changing environmental conditions.

By modelling genomic variation across the landscape, we can understand the environmental factors that shape patterns of genomic variation and identify variation suitable for predicted future climates. We found several environmental variables that have played a role in the structure of genomic variation across the landscape. Of these, the climate variables are predicted to change rapidly over time. Change in soil nitrogen content might occur over longer time scales, but it is difficult to forecast due to complex biotic feedbacks (Brevik, 2013). This suggests that optimal seed sourcing will need to balance the tracking of rapidly changing climate variables with the need to account for variables that are more stable due to their dependence on stable features of geology, topography, or hydrology. This also highlights an important concern that key environmental variables may become uncoupled, resulting in less than ideal conditions for this species across the landscape.

Our analyses of phenotypic variation found no site-specific variation in phenotypic plasticity that would enable us to identify provenances better able to cope with rapid environmental change. However, plasticity is trait specific and traits that are hypothesized to be important for establishment and survival should continue to be investigated because they may provide valuable information for restoration projects. Importantly, our growth experiments support the results of the genomic analyses, showing the remarkable extent of variation both among sites and within sites, further supporting our recommendation to source seed to incorporate the high level of variation that occur naturally in *E. melliodora*.

The results of this study are promising for the future of *E. melliodora* across its native distribution. We found high genomic and phenotypic diversity within sites, as well as shared across the range. This naturally occurring variation can provide a basis for adaptation to a rapidly changing environment and it should be incorporated into restoration projects through strategic seed sourcing. It is important to note that our genomic analyses were based on mature trees that predate extensive land clearing for agriculture. The same analyses in seedlings or saplings at these sites may show different results, although our phenotypic studies using seedlings produced concordant results. It remains to be determined if human modifications of the landscape have disrupted historical patterns of gene flow, resulting in more fragmented and inbred populations.

Our landscape genomic model can guide seed selection by empirically defining local provenances and identifying variation suitable for predicted future climates. This understanding of the relationship between environmental and genomic variation can be combined with other types of information, such as basic biological knowledge of the ecological community and best agronomic practices in restoration, to establish foundation species and ecosystems with the highest probability of success in a rapidly changing environment.

## Methods

### Sample Collection

We obtained *E. melliodora* leaf samples from mature trees at 38 sites across the species’ range through a community science project described in Broadhurst et al. (*in review*) (Supplemental Table S1). From each site, a citizen scientist collected leaf samples from up to 30 trees, put the samples in silica gel for drying, and shipped them to CSIRO for processing. In addition to leaf material, they also collected seeds from the sampled trees when available. We sampled an additional seven trees planted at a single site in Western Australia, well outside the species’ natural distribution.

### Genotyping by Sequencing

We selected 3 to 10 trees per sampling site for sequencing and we processed each of the seven trees from Western Australia twice, using different leaves from the same tree, to serve as technical replicates. No power analysis was used to determine sample size during the design of the study. Sample size was determined based on our experience and judgment, with consideration of the availability of samples. We sequenced these 379 samples using a modified Genotyping-By-Sequencing (GBS) protocol (Elshire et al., 2011). Briefly, we extracted genomic DNA from approximately 50 mg of leaf tissue using the Qiagen DNeasy Plant 96 Kit, digested with PstI for genome complexity reduction, and ligated with a uniquely barcoded sequencing adapter pair. We then individually PCR amplified each sample to avoid sample bias. We pooled samples in equimolar concentrations and extracted library amplicons between 350 and 600 bp from an agarose gel. We sequenced the library pool on an Illumina HiSeq2500 using a 101- bp paired-end protocol at the Biomolecular Resource Facility at the Australian National University, generating almost 260 million read pairs.

We checked the quality of the raw sequencing reads with FastQC (v0.10.1, (Andrews, 2012)). We used AXE (v0.2.6, (Murray & Borevitz, 2017a)) to demultiplex the sequencing reads according to each sample’s unique combinatorial barcode and were unable to assign 11% of read pairs to a sample. We used trimit from libqcpp (v0.2.5, (Murray & Borevitz, 2017b)) to clean the reads for each sample, using default parameters, except q=20. This involved removing adapter contamination due to read-through of small fragments (20% of read pairs) and merging overlapping pairs (49% of read pairs), both steps using algorithms based on a global alignment of read pairs. We also used trimit for sliding window quality trimming (11% of reads). We then used a custom script to remove sequencing reads that did not begin with the expected restriction site sequence (16% of reads). We aligned sequencing reads to the *E. grandis* reference genome (v2.0, (Bartholomé et al., 2015; JGI, 2015; Myburg et al., 2014)), including all nuclear, chloroplast, mitochondrial, and ribosomal scaffolds, but used only nuclear scaffolds for downstream analyses. We aligned reads using bwa-mem (v0.7.5a-r405, (Li, 2013)), as paired reads (-p) and treating shorter split hits as secondary alignments (-M), with 88% of reads successfully mapped. We used GATK’s HaplotypeCaller in GVCF mode (v3.6-0-g89b7209, (McKenna et al., 2010)) to call variants for each sample with heterozygosity (-hets) increased to 0.005, indel heterozygosity (-indelHeterozygosity) increased to 0.0005, and the minimum number of reads sharing the same alignment start (-minReadsPerAlignStart) decreased to 4.

We used GATK’s GenotypeGVCFs (v3.6-0-g89b7209, (McKenna et al., 2010)) for a preliminary round of joint genotyping across all samples based on the individual variant calls and again increasing the heterozygosity (-hets) to 0.005 and the indel heterozygosity (-indelHeterozygosity) to 0.0005. For basic filtering, we used GATK to remove variants that were indels, had no variation relative to the reference, were non-biallelic SNPs, had QD<2.0 ("variant call confidence normalized by depth of sample reads supporting a variant"), MQ>40.0 ("Root Mean Square of the mapping quality of reads across all samples"), or MQRankSum<-12.5 ("Rank Sum Test for mapping qualities of REF versus ALT reads"). We removed samples with more than 60% missing data and SNPs with more than 80% missing data. We examined the genomic distance between samples to verify that technical replicates clustered closely together. We used a preliminary PCA, based on genomic distance between samples, to identify outlier samples. We removed outlier samples and poorly sequenced samples from the dataset for final genotyping and all downstream analyses.

We reran GATK’s joint genotyping on the final sample set. We again used GATK to remove variants that were indels, SNPs with no variation relative to the reference, and non-biallelic SNPs. We determined final filtering thresholds by examining parameter distributions. A locus was retained for subsequent analysis if ExcessHet<13.0 ("phred-scaled p-value for exact test of excess heterozygosity"),-0.3<InbreedingCoeff<0.3("likelihood-based test for the inbreeding among samples"), MQ>15.0 ("Root Mean Square of the mapping quality of reads across all samples"),-10.0<MQRankSum<10.0 ("Rank Sum Test for mapping qualities of REF versus ALT reads"), and QD>8.0 ("variant call confidence normalized by depth of sample reads supporting a variant"). We ran a second preliminary PCA analysis to identify additional outlier samples. Finally, we used VCFtools (v0.1.12b, (Danecek et al., 2011)) to remove SNPs with greater than 60% missing data and thin the SNPs so that none were closer than 300 bp.

### Genomic Analyses

To examine the genomic structure of *E. melliodora* and how it is influenced by geography, we conducted individual-based analyses. For these analyses, we converted the final genotypic data (a vcf file) to a sample-by-SNP matrix and imported it into a genind object (R adegenet v2.0.1, (Jombart, 2008)). We calculated the pairwise genomic distances between individuals using a euclidean distance in *dist* (R stats v3.1.2, (R Core Team, 2015)). To visualize the genomic distance among samples, we ran a PCA using *dudi.pco* (R ade4 v1.7-4, (Dray & Dufour, 2007)). We plotted the first two PCA components, with samples colored in a rainbow gradient based on sample latitude. We calculated the geographic distance between samples based on their GPS coordinates using *earth.dist* (R fossil v0.3.7, (Vavrek, 2011)). We used a *mantel* test (R vegan v2.4-0, (Oksanen et al., 2016)) to quantify the linear relationship between the genomic distance between individuals and the natural log of geographic distance.

To examine the role that environmental factors play in driving the genomic structure across the landscape, we used Generalized Dissimilarity Modelling (GDM), which uses matrix regression to estimate the non-linear relationship between genomic distance and environmental distance (Ferrier et al., 2007; Fitzpatrick & Keller, 2015; Thomassen et al., 2011). We then used this model to predict the distribution of genomic variation across the landscape under current environmental conditions, as well as predicted future conditions. We obtained environmental data for the GDM from climate, elevation, soil, and landscape raster layers. Climate variables included 19 bioclimatic variables for the current time period (1960-1990), at 30 arc second resolution (WorldClim, 2016b). Elevation was from a digital elevation model aggregated from 90 m resolution (CGIAR-CSI, 2016). Soil data included available water capacity, total nitrogen, and total phosphorus sampled at the surface (0-5 cm) and at depth (100-200 cm) (CSIRO, 2016). Landscape data included the Prescott Index (a measure of water balance) and MrVBF (a topographic index) (CSIRO, 2016). For future prediction, we used the 19 bioclimatic variables predicted for 2070 at 30 arc second resolution based on GCM MIROC5 for representative concentration pathway 8.5 (WorldClim, 2016a), which is a greenhouse gas concentration trajectory showing continual increase in emissions over time. We determined the values for each variable at each sampling site based on GPS coordinates and used those values to calculate the environmental distances between sites.

To determine the genomic distances between sampling sites, we used the sample by SNP matrix to calculate pairwise F_st_ (Weir & Cockerham, 1984) using *pairwise.WCfst* (R hierfstat v0.04-22, (Goudet & Jombart, 2015)). We ran a sampling site level PCA on the pairwise F_st_ matrix using *dudi.pco* (R ade4 v1.7-4, (Dray & Dufour, 2007)) and calculated the percent of variation explained for each PCA axis. For the GDM, we scaled the F_st_ matrix to between 0 and 1 by subtracting the minimum value and then dividing by the maximum value. We generated the GDM model using *gdm* (R gdm v1.2.3, (Manion et al., 2016)) with the scaled F_st_ matrix, geographic distance between sites, and environmental distances for the 28 variables for the current time period. Initially, we generated a GDM model for each environmental variable separately and excluded variables from further analysis if the deviance explained by the model was less than 5%. For the remaining variables, we calculated Pearson’s correlation for site values between pairwise sets of variables. If a pair of variables had a correlation greater than 60%, we excluded the variable with the lowest explanatory power from subsequent analysis. We conducted permutation testing using gdm.varImp (R gdm v1.2.3, (Manion et al., 2016)) with 1000 permutations to determine the explanatory power and statistical significance of the remaining variables and excluded additional inconsequential variables. We generated a final GDM model with the remaining environmental variables.

We cross validated the GDM model using a random 70% of the spatial sampling sites as training data and the remaining 30% of sites as test data and ran 1000 resampled iterations. We used the GDM models from the training sites to predict the genomic dissimilarity between the test sites and used Pearson’s correlation to compare the predicted values to the observed values. To test the robustness of the geographic prediction from the GDM model, we visualized the geographic splines from 100 of these GDM models.

To project the final GDM model onto the current environmental landscape, we first delineated the geographic extent of the analysis by defining an *E. melliodora* distribution polygon. We downloaded 14,977 *E. melliodora* occurrence records from the Atlas of Living Australia (ALA, 2016), of which we removed 189 because they were well outside the expected distribution or were sparse records on the distribution margin. We generated the polygon using ahull (R alphahull v2.1, (Pateiro-López & Rodríguez-Casal, 2010)), with alpha=15 and gBuffer (R rgeos v0.3-21, (Bivand & Rundel, 2016)), with a 20 km buffer. We then transformed the environmental rasters based on the model splines (*gdm.transform*), took a PCA of the transformed layers (*prcomp* R stats v3.1.2, (R Core Team, 2015)), and predicted across space (*predict*). We visualized the result by graphing the first three components of a PCA using a red-green-blue plot (Fitzpatrick & Keller, 2015). We also projected the model onto a predicted future environmental landscape with the same procedure, except we substituted the current bioclimatic rasters with the future ones for 2070 that were predicted under a high CO2 emission scenario. We calculated the expected change in the distribution of genomic variation over time using the *predict* function with time=T.

We examined the implications of the GDM model for seed sourcing decisions by selecting two hypothetical reforestation sites. We compared predicted future GDM values at these two hypothetical reforestation sites to current climate GDM values across the landscape of potential seed sources. This enabled us to generate a map of predicted genomic similarity of potential seed sources to the hypothetical reforestation sites under climate change.

### Growth Experiments

To examine the effect of provenance and environment on phenotype, we conducted experiments in climate controlled growth chambers under two different climate regimes. No power analysis was used to determine sample size during the design of the experiment. Sample size was determined based on our experience and judgment, with consideration of the availability of seed and space in the growth chambers. We selected six sites (11, B, D, G, T1, T3) and six maternal trees per site that had sufficient seed (asterisks in Fig. 1A). For each chamber, we grew eight or nine replicate seedlings from each maternal tree. To ensure we had a seedling for each intended replicate, four seeds were planted per pot (6.5 cm x 6.5 cm x 20 cm pots with soil that was 80% Martin’s mix and 20% sand). We germinated seeds in climate controlled chambers with 12 hours of light at 25°C and 12 hours of dark at 15°C. We set lights to mimic summer morning light (photosynthetic photon flux 370 nm=82, 400 nm=83, 420 nm=78, 450 nm=37, 530 nm=31, 620 nm=72, 660 nm=28, 735 nm=34, 850 nm=89, 6500 K=94 μmol/m^2^/s). We watered all seeds twice daily to keep the soil moist. We culled to one seedling per pot 12-14 days after planting.

Three weeks after germination, we sorted seedlings into treatment chambers based on a randomized block design. Climate conditions were determined with SolarCalc (Spokas & Forcella, 2006) to mimic average summer conditions (sampling site 11) and hotter conditions (5°C temperature increase; sampling site T3). We ran the experimental conditions for 12-14 weeks and took phenotypic measurements at five time points (1, 2, 3, 5, and 11 weeks after the experimental treatment began). Measurements included seedling height, number of leaves, and total leaf length.

For the analysis of seedling height and total leaf length, we used the measurements at five weeks after the experimental treatment began and used only seedlings that were determined to be well established at that time. We also calculated a relative height increment for each seedling by determining the last measurement when the seedling had two or fewer leaves and the first measurement with eight or more leaves. The relative height increment is the difference between the natural log of the two height measures, divided by the difference in time.

We investigated phenotypic plasticity by examining interaction plots between maternal line and experimental conditions for three response variables: seedling height, total leaf length, and relative height increment. We statistically tested for an interaction between sampling site/maternal line and experimental condition with linear mixed-effect models using *lmer* (R lme4 v1.1-10, (Bates et al., 2015)) for each of the three response variables. Due to a lack of power to consider maternal line nested within sampling site, we ran two models for each response variable—one with maternal line and one with sampling site. These models included the experimental condition, sampling site or maternal line, and and their interactions as fixed effects. We included germination chamber and block as random effects. We identified outliers visually and ran the models with and without outliers to determine if they affected the results.

We visualized the distribution of values for the three response variables across the six sampling sites using box plots. We quantified the distribution of phenotypic variation with linear mixed-effect models using *lmer* (R lme4 v1.1-10, (Bates et al., 2015)). For each of the three response variables, the model included maternal line nested within sampling site and experimental condition as main effects, with no interaction term, and germination chamber and block as random effects.

After completion of the chamber experiment, we conducted an outdoor covered drought experiment on the 16 week old seedlings. No power analysis was used to determine sample size during the design of the experiment. Sample size was determined based on our experience and judgment, with consideration of the availability of space in the covered growth facility. We selected 160 seedlings from five sampling sites, with four maternal lines per site. We paired each seedling with a seedling of similar size from the same maternal line and treatment chamber. We randomly assigned each seedling of the pair to a different drought treatment group. We transplanted the seedlings to PVC tubes (9 cm diameter x 50 cm height with sand, perlite, and slow release osmocote) and kept them well watered for seven weeks, allowing them to acclimate to the outdoor conditions. Then we imposed two treatments: well watered and drought. For the well watered treatment, we watered the seedlings to saturation as needed (between three times per week and twice per day, depending on the weather). For the drought treatment, we watered as necessary to reach (but not exceed) 50% saturation.

We measured leaf traits on each seedling three weeks after the initiation of treatment. We measured stomatal conductance with a porometer (SC-1 Leaf Porometer by Decagon Devices) and determined that water stress was induced in the droughted seedlings. We determined the leaf length to width ratio from a scan of the most recent fully expanded leaf from each seedling using image analysis software (WD3 WinDIAS Leaf Image Analysis System by Delta-T Devices). This leaf was initiated prior to the start of treatment, but expanded while under treatment conditions. We took additional measurements two months after the initiation of treatment. We used a chlorophyll meter (SPAD – 502 by Konica Minolta) to determine the SPAD index, which measures relative chlorophyll content; reduction in SPAD index would indicate detrimental effects of water limitation. We calculated specific leaf area (SLA, leaf area divided by dry mass) by scanning a single leaf from each seedling to determine the leaf area (WD3 WinDIAS Leaf Image Analysis System by Delta-T Devices) and weighed oven dried leaves. For analysis, we excluded data for seedlings that died during the experiment. We also excluded the experimental treatment pair of any dead seedlings.

We visualized phenotypic plasticity by examining interaction plots between maternal line and experimental conditions for four response variables: stomatal conductance, leaf length to width ratio, SPAD index, and SLA. We statistically tested for an interaction between sampling site/maternal line and experimental condition with linear mixed-effect models using *lmer* (R lme4 v1.1-10, (Bates et al., 2015)) for each of the four response variables. Due to a lack of power to consider maternal line nested within sampling site, we ran two models for each response variable-one with maternal line and one with sampling site. These models included the experimental condition, sampling site or maternal line, and their interactions as fixed effects. We included block and sample pairings as random effects.

We visualized the distribution of values for the four response variables across the five sampling sites using box plots. We quantified the distribution of phenotypic variation with linear mixed-effect models using *lmer* (R lme4 v1.1-10, (Bates et al., 2015)). For each of the four response variables, the model included maternal line nested within sampling site and experimental condition as main effects, with no interaction term, and block and sample pairings as random effects. Due to a lack of power, the p-value for the sampling site term was determined from a model without maternal line.

## Acknowledgments

We thank the ANU Bioinformatics Consultancy for computational support and advice, the ANU Statistical Consulting Unit for statistical advice, the Centre for Biodiversity Analysis for advice on GDM modelling, and the ANU NCRIS Plant Growth Facility and RSB Plant Services for assistance with growth experiments.

## Data Access

GBS sequencing reads are available at the NCBI Sequence Read Archive (SRA) (http://www.ncbi.nlm.nih.gov/sra) under BioProject PRJNA413429. Growth experiment data and scripts for genomic and phenotypic analyses are available at https://github.com/LaMariposa/emelliodora.

## Supplementary Figures

**Figure S1.**
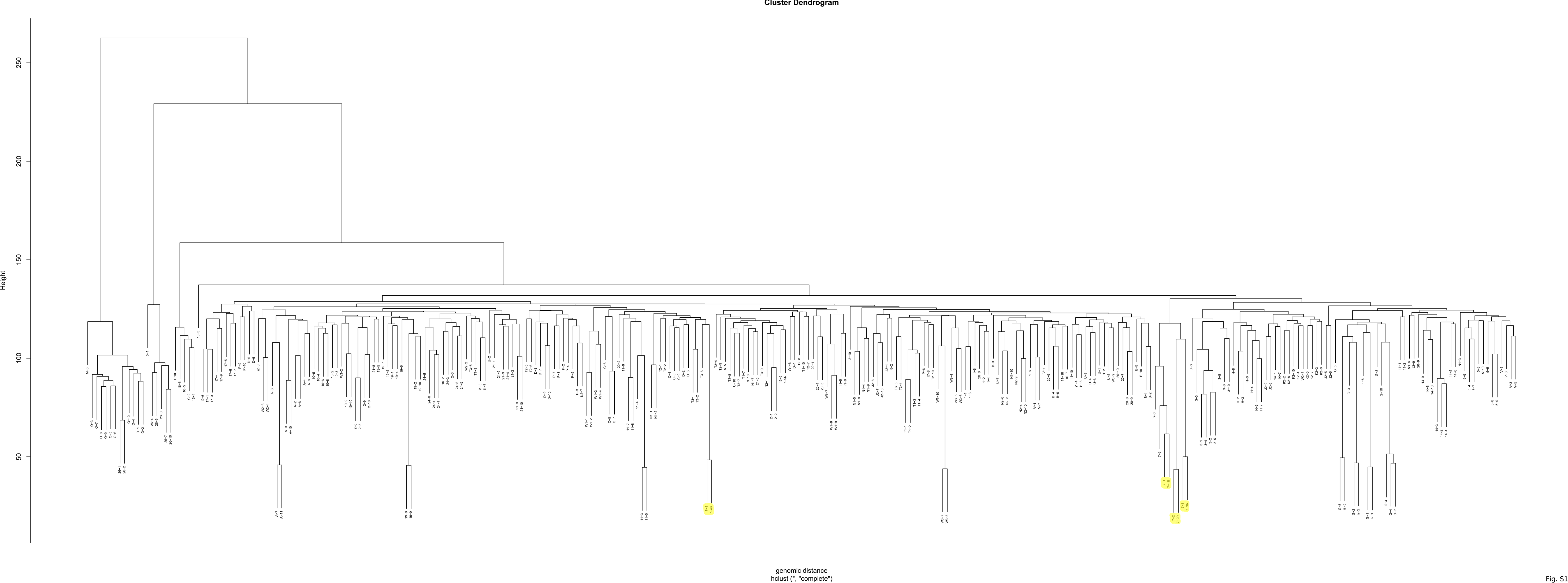
Technical replicate dendrogram. Dendrogram based on genomic distance between samples showing the strong clustering of technical replicates (denoted with an "R" after the sample name and highlighted in yellow). Note that three of the technical replicates failed to pass quality control and are not included in the dendrogram. Additional sample pairs show strong clustering. In each case, the individuals of the pair are from the same sampling site, indicating samples that are closely related.

**Figure S2.**
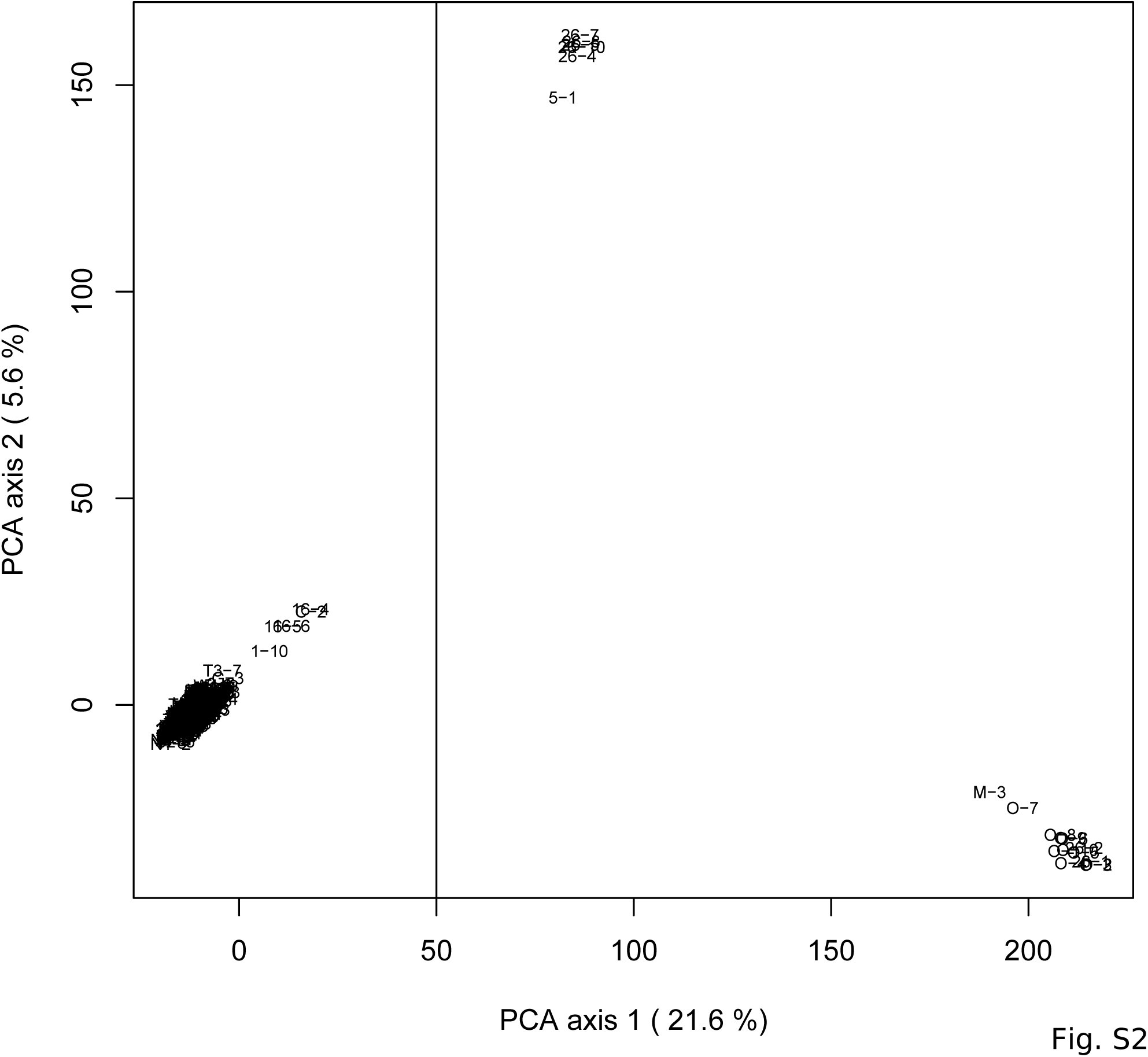
Species identification PCA. PCA of genomic distance between samples showing strong outliers that are likely misidentified samples or hybrids. The vertical line at 50 on PCA axis 1 indicates the cutoff, with all samples to the right removed from further analyses.

**Figure S3.**
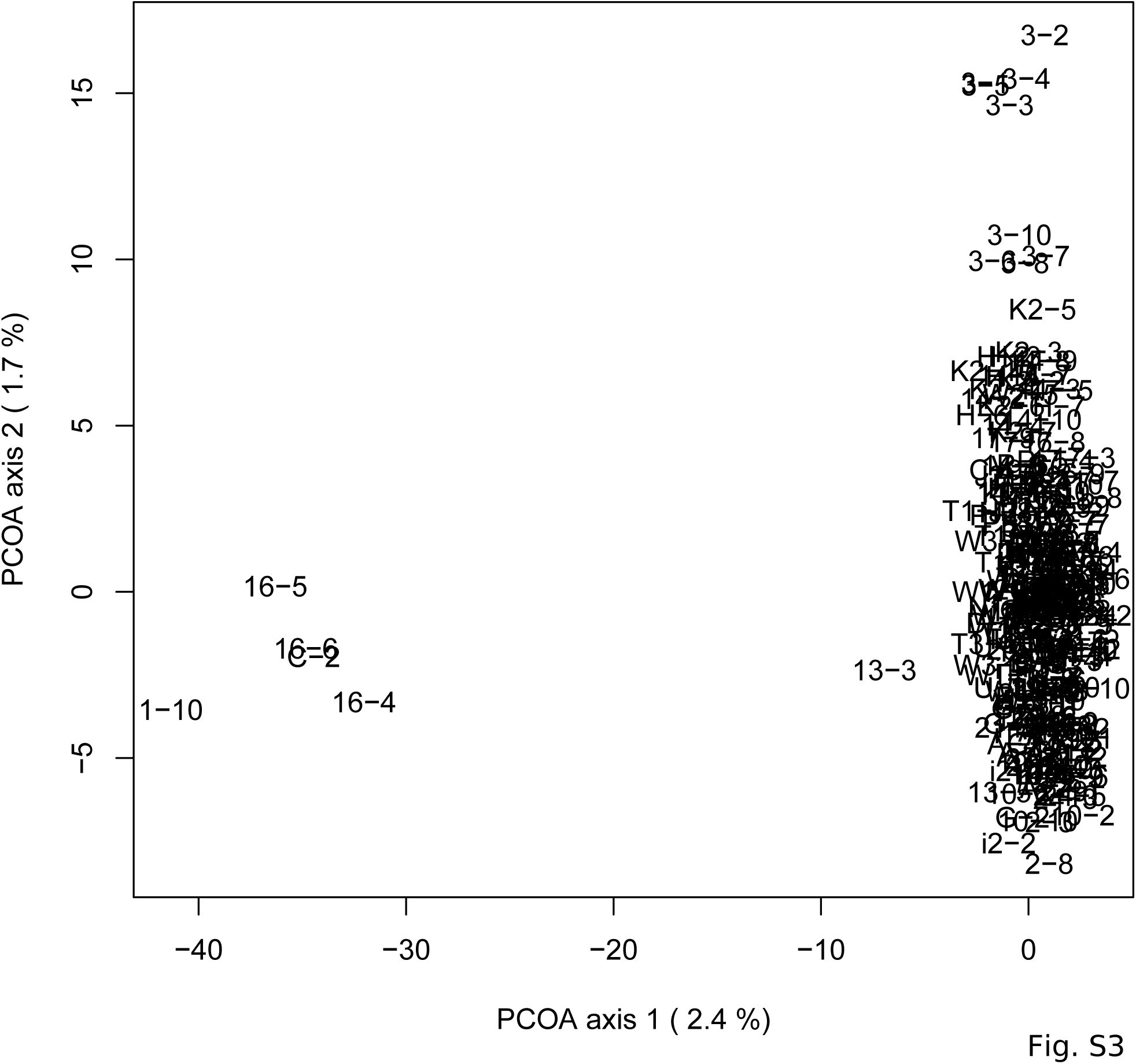
Outlier PCA. PCA of genomic distance between samples for the confirmed *E. melliodora* samples. The five samples on the left were deemed outliers and removed from further analyses.

**Figure S4.**
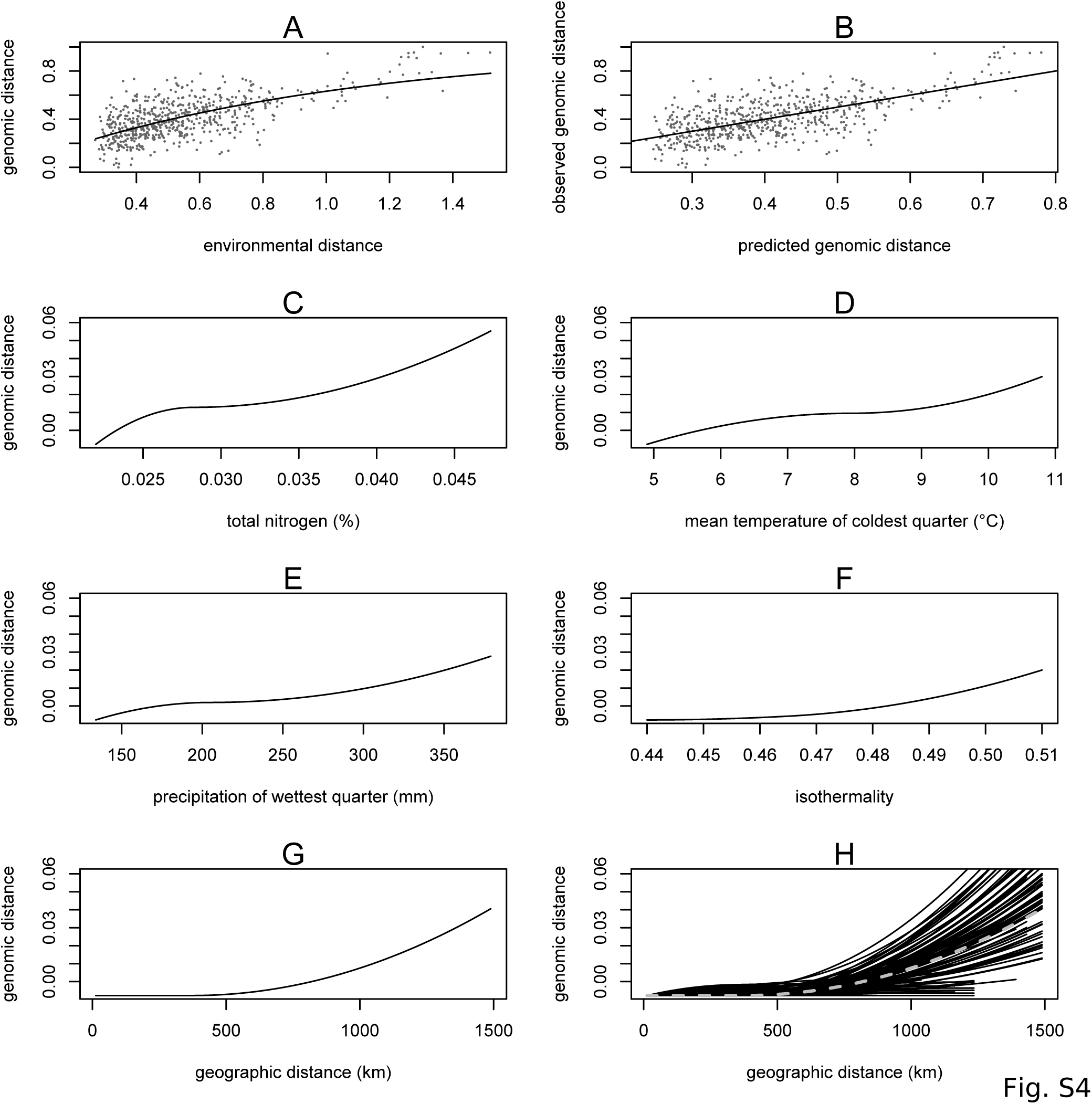
Generalized dissimilarity modelling (GDM) results. (A) Non-linear relationship between environmental distance and genomic distance. Points are site pairs; the line is the predicted relationship. (B) Relationship between predicted genomic distance and observed genomic distance. Points are site pairs; the line indicates where observation and prediction match. (C-G) Predicted splines showing the estimated relationship between the environmental variable and genomic distance for (C) total nitrogen content at 100- 200 cm of soil depth, (D) mean temperature of the coldest quarter, (E) precipitation of the wettest quarter, (F) isothermality, and (G) geography. (H) Geographic splines from 100 iterations of sampling 70% of sites. Each solid black line is an iteration; dashed grey line is the full model prediction.

**Figure S5.**
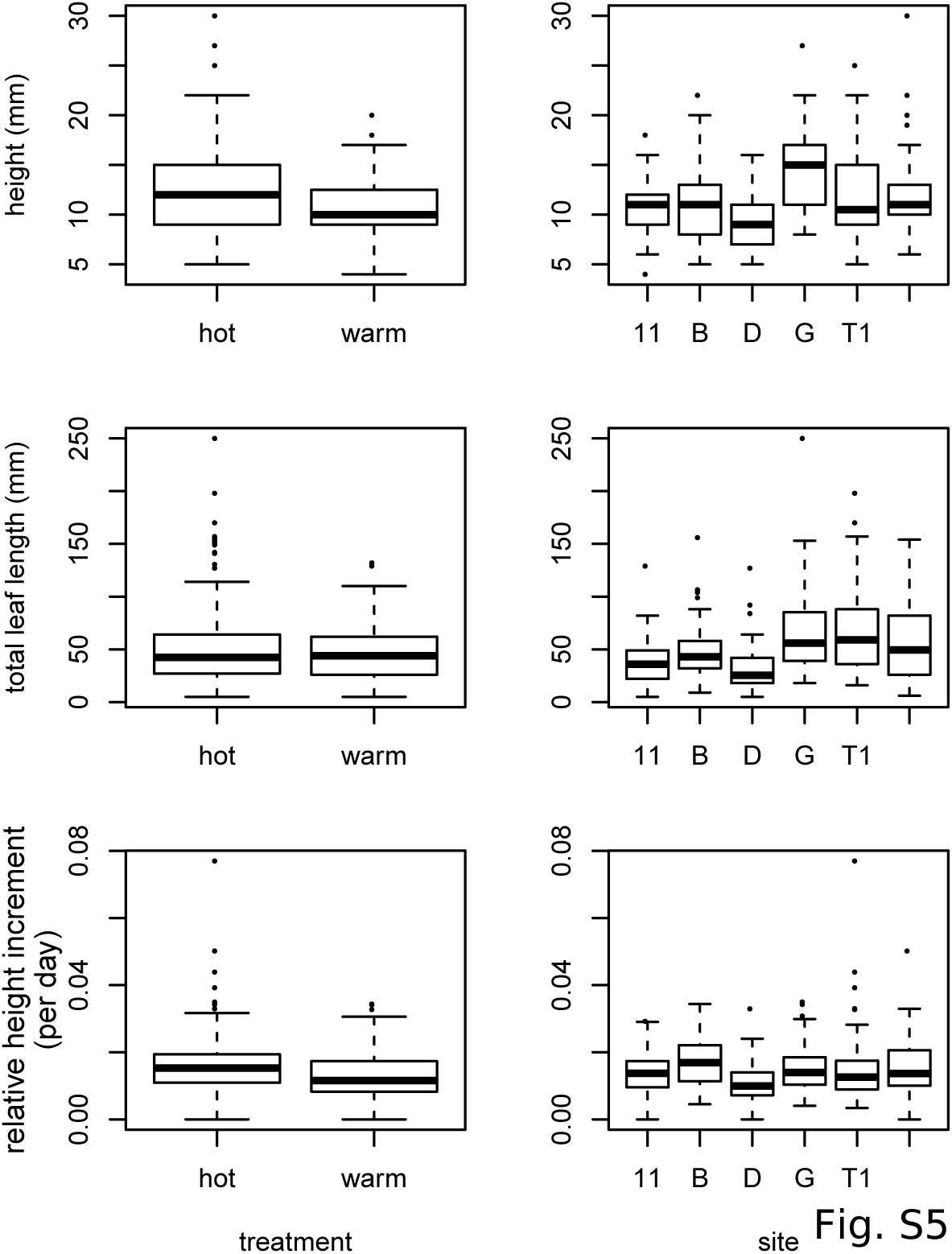
Variation in seedling growth in chamber experiment. Box plots showing variation between chamber treatments (left) and sampling sites (right) for three seedling growth traits.

**Figure S6.**
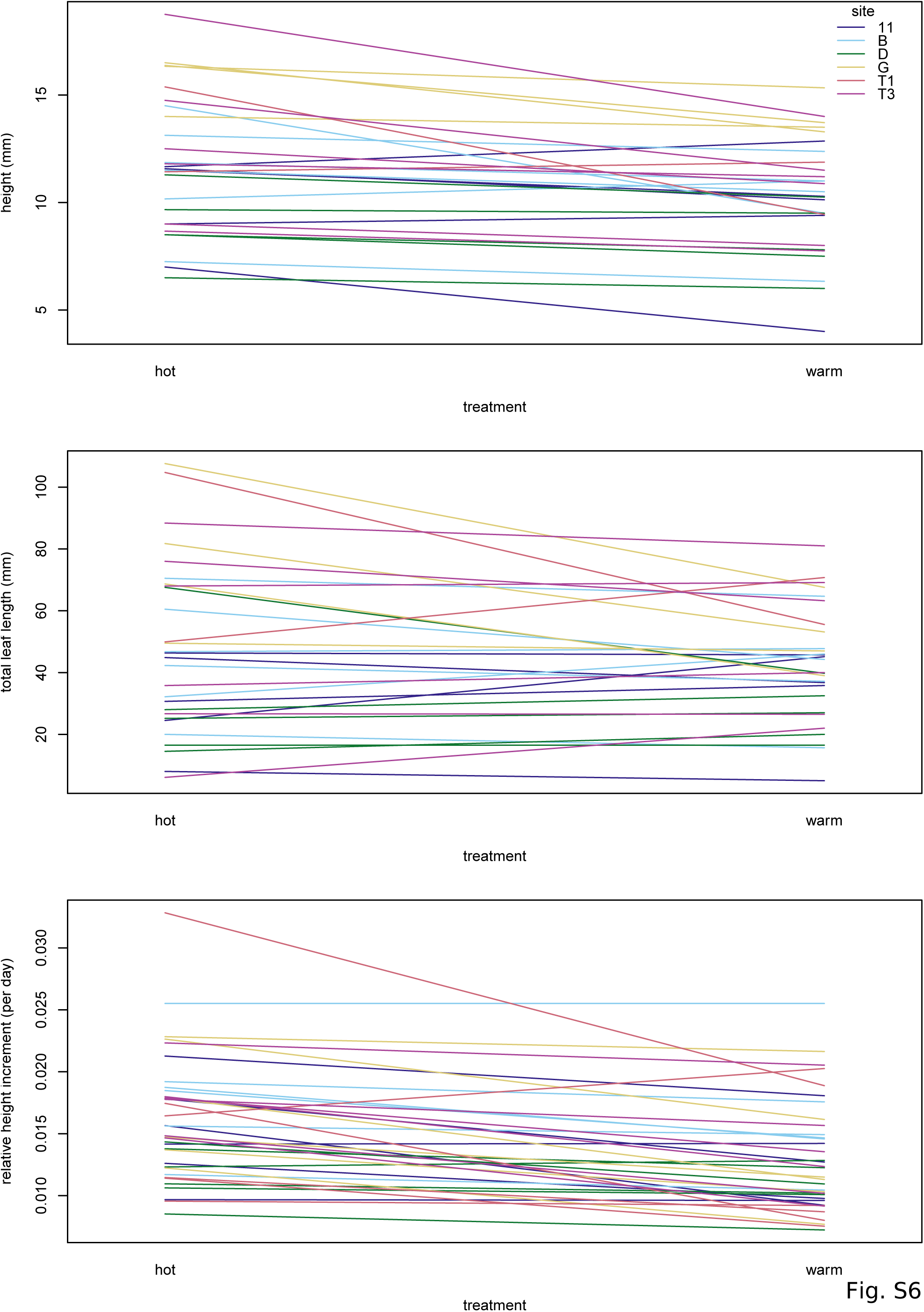
Interaction plots for chamber experiment. Plots showing interactions between three seedling growth traits and the experimental conditions. Each line represents a maternal line, with color indicating the sampling site.

**Figure S7.**
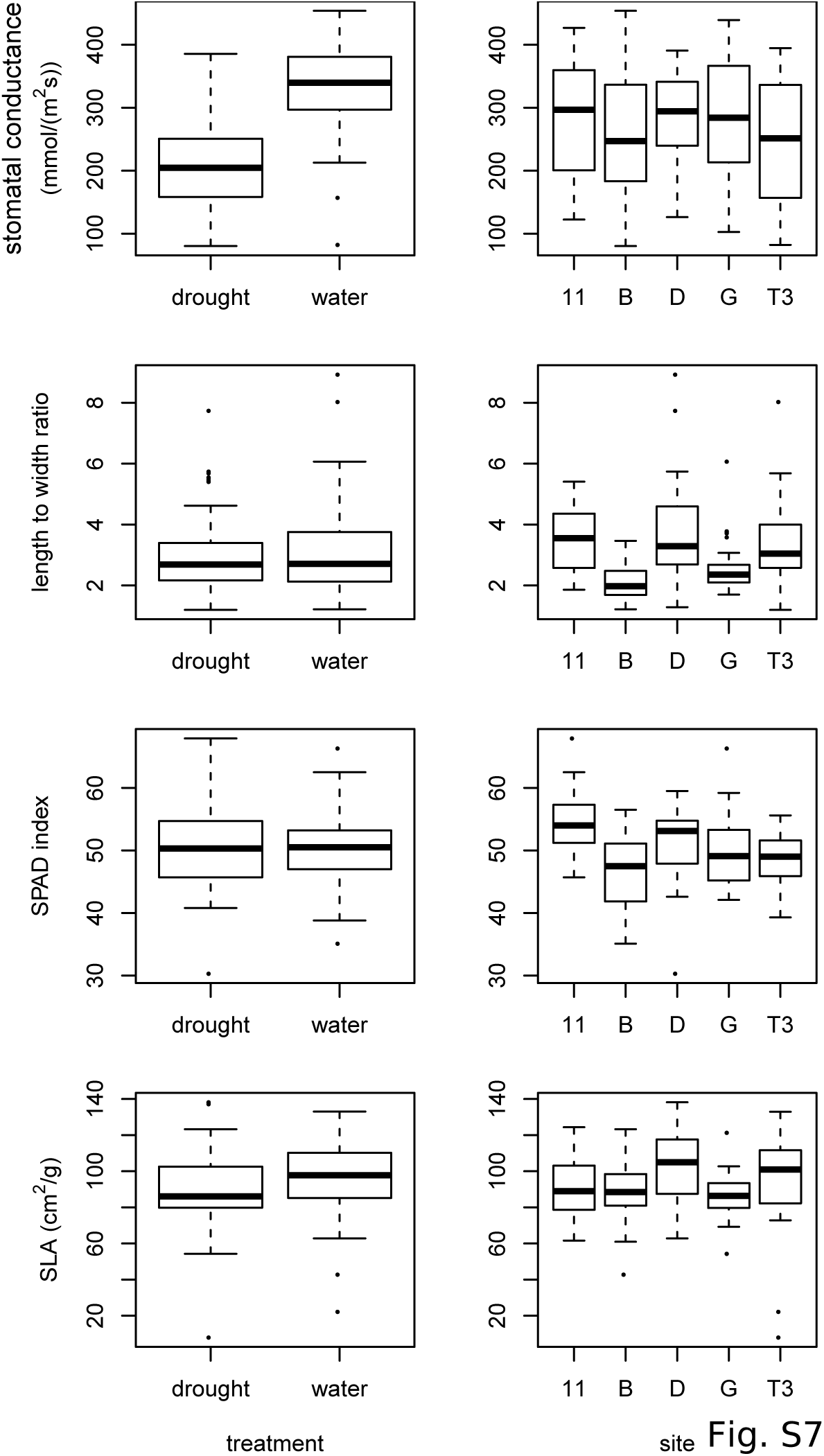
Variation in leaf traits in drought experiment. Box plots showing variation between water treatments (left) and sampling sites (right) for four leaf traits.

**Figure S8.**
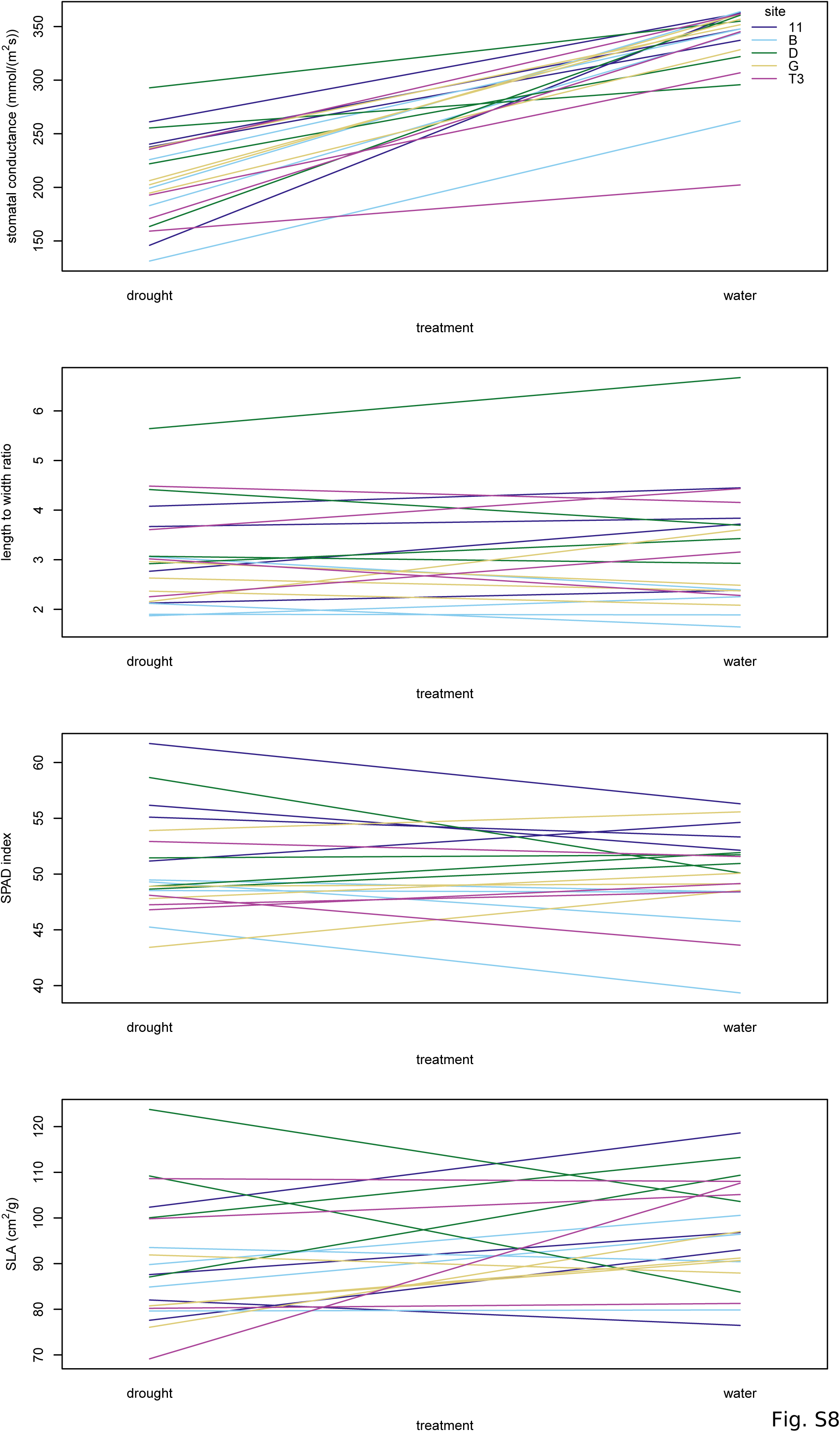
Interaction plots for drought experiment. Plots showing interactions between the four leaf traits and the water treatment. Each line represents a maternal line, with color indicating the sampling site.

**Figure S9.**
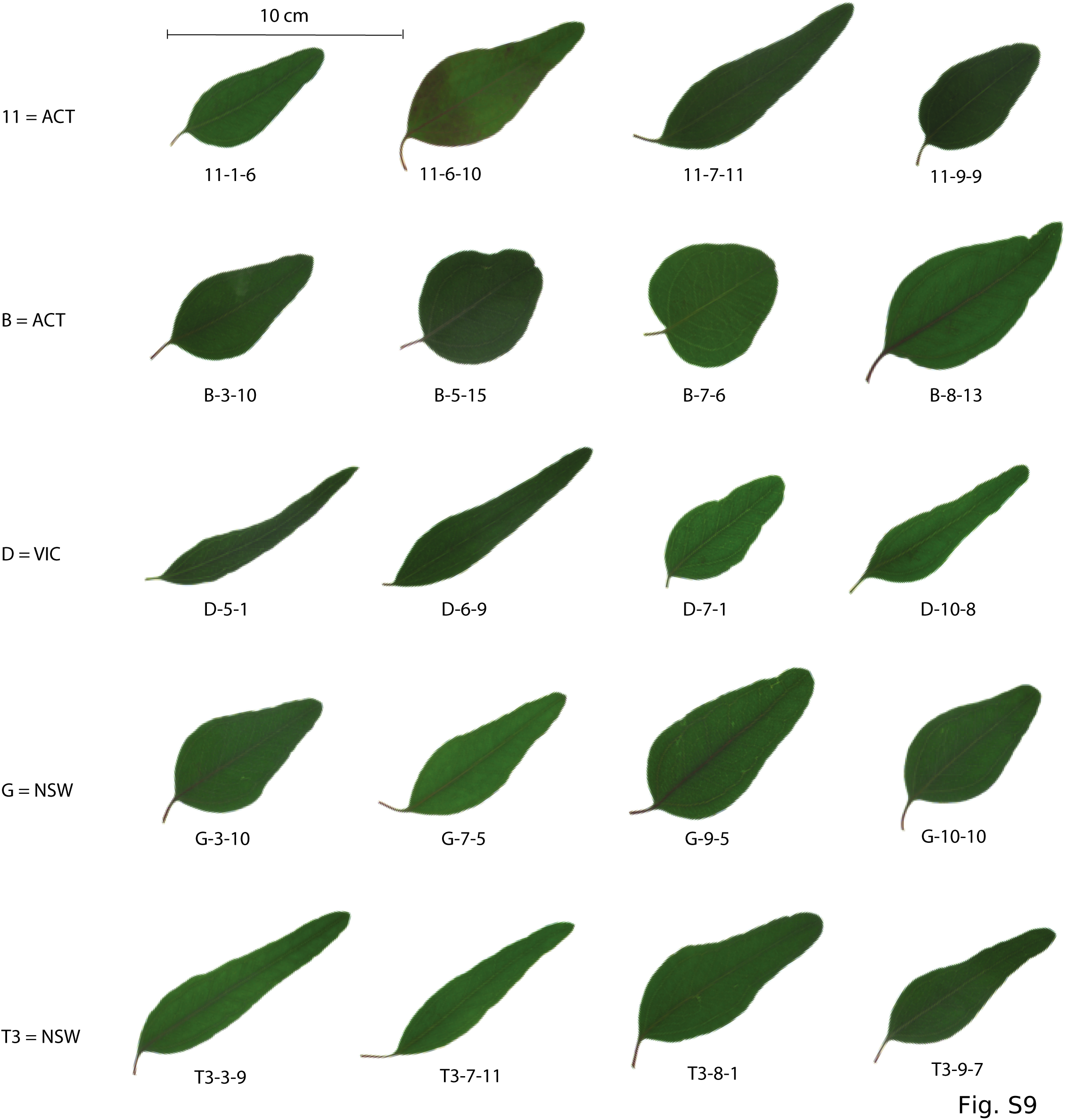
Variation in leaf shape. One representative leaf from each maternal line in the drought experiment. Each row shows a single sampling site, identified by site ID and state location (ACT=Australian Capital Territory, VIC=Victoria, NSW=New South Wales). Each leaf is identified by its sampling site, maternal line, and replicate number).

**Table S1:** *E. melliodora* sampling information

| site name | latitude | longitude | # sampled | # outliers* | # in final analysis | chamber experiment | drought experiment |
| --- | --- | --- | --- | --- | --- | --- | --- |
| 1 | -33.416668 | 149.55055 | 10 | 1 secondary | 7 |  |  |
| 2 | -36.6755 | 146.26477 | 10 |  | 7 |  |  |
| 3 | -27.368696 | 152.03023 | 10 |  | 9 |  |  |
| 5 | -33.710145 | 149.52355 | 10 | 1 primary | 9 |  |  |
| 7 | -33.936944 | 116.963333 | 7 | 7 geographic | 0 |  |  |
| 10 | -37.285278 | 143.77778 | 10 |  | 8 |  |  |
| 11 | -35.40026 | 149.1318 | 10 |  | 9 | yes | yes |
| 13 | -36.997778 | 145.474944 | 6 |  | 3 |  |  |
| 14 | -32.389167 | 150.881667 | 10 |  | 10 |  |  |
| 16 | -33.224724 | 149.28056 | 7 | 3 secondary | 2 |  |  |
| 17 | -33.44218 | 147.51628 | 10 |  | 8 |  |  |
| 19 | -37.179444 | 144.483333 | 10 |  | 9 |  |  |
| 20 | -34.911785 | 148.971179 | 10 |  | 9 |  |  |
| 21 | -37.144361 | 142.839428 | 10 |  | 9 |  |  |
| 24 | -36.980278 | 144.047611 | 10 |  | 7 |  |  |
| 26 | -35.39474 | 149.677129 | 10 | 7 primary | 2 |  |  |
| A | -36.388 | 146.481 | 10 |  | 10 |  |  |
| B | -35.235664 | 149.16422 | 10 |  | 7 | yes | yes |
| C | -35.03333 | 147.33333 | 9 | 1 secondary | 6 |  |  |
| D | -36.3675 | 145.70232 | 10 |  | 9 | yes | yes |
| G | -36.86476 | 148.85016 | 10 |  | 9 | yes | yes |
| H | -36.63518 | 149.56728 | 10 |  | 7 |  |  |
| i1 | -36.75 | 145.583333 | 10 |  | 8 |  |  |
| i2 | -36.54041 | 146.10739 | 10 |  | 7 |  |  |
| J2 | -33.121143 | 149.05785 | 10 |  | 9 |  |  |
| K2 | -30.54493 | 151.79367 | 10 |  | 8 |  |  |
| M | -34.630472 | 149.870892 | 3 | 1 primary | 0 |  |  |
| N1 | -34.036884 | 148.5675 | 10 |  | 10 |  |  |
| N2 | -33.6166 | 149.64452 | 10 |  | 7 |  |  |
| O | -37.4279 | 147.8924 | 10 | 10 primary | 10 |  |  |
| P | -35.819633 | 145.31187 | 10 |  | 9 |  |  |
| T1 | -35.26495 | 147.310133 | 10 |  | 7 | yes |  |
| T2 | -35.37515 | 147.25261 | 10 |  | 9 |  |  |
| T3 | -35.299683 | 147.165733 | 10 |  | 10 | yes | yes |
| U | -36.632668 | 144.35019 | 10 |  | 8 |  |  |
| V | -33.423 | 149.752 | 10 |  | 7 |  |  |
| W1 | -36.42797 | 144.41092 | 10 |  | 8 |  |  |
| W2 | -36.580471 | 143.521639 | 10 |  | 4 |  |  |
| W3 | -36.980278 | 144.047611 | 10 |  | 8 |  |  |

\* primary outliers refers to outlier samples identified with the first PCA analysis secondary outliers refers to outlier samples identified with the second PCA analysis geographic refers to samples outside the natural distribution

**Table S2:**
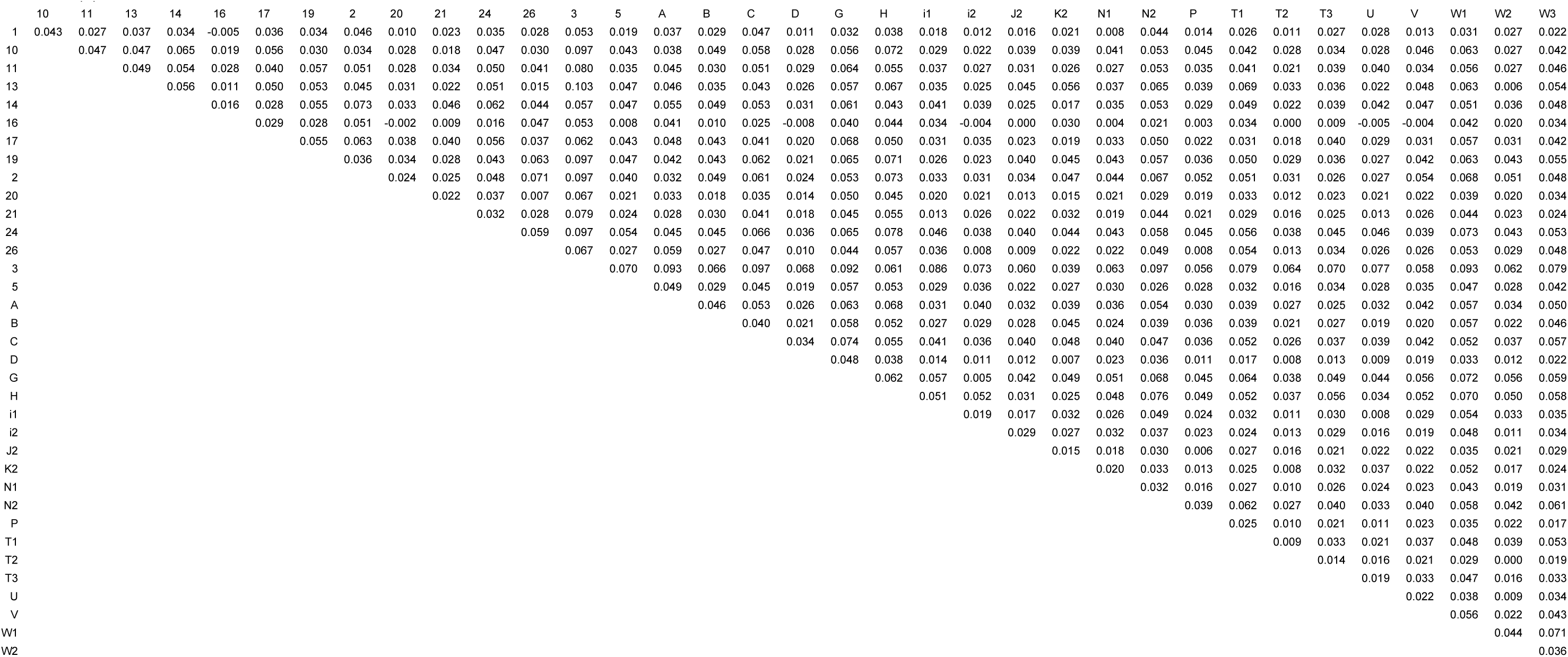
Pairwise population Fst

**Table S3:**
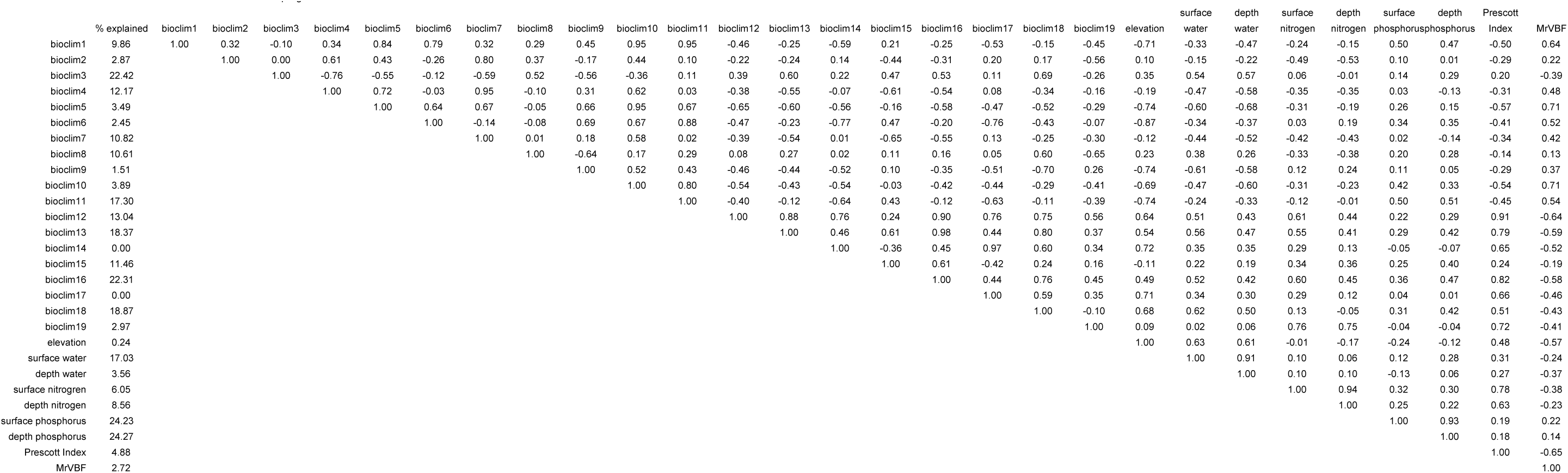
Pearson’s correlation between environmental variables across sampling sites

**Table S4:** Percent of variation explained and p-values for non-interaction linear models for chamber experiment

|  | seedling height |  | total leaf length |  | relative height increment |  |
| --- | --- | --- | --- | --- | --- | --- |
|  | % explained | p-value | % explained | p-value | % explained | p-value |
| sampling site | 17.7 | <0.00001 | 8.2 | 0.00063 | 1.8 | <0.00001 |
| maternal line:sampling site | 10.6 | 0.00031 | 17.2 | 0.00001 | 27.6 | <0.00001 |
| experimental condition | 5.4 | 0.00032 | 1.2 | 0.05084 | 8.1 | <0.00001 |
| germination chamber | 0 | -- | 0.3 | -- | 0.8 | -- |
| block | 1.1 | -- | 1.6 | -- | 5.1 | -- |
| residual | 65.2 | -- | 71.5 | -- | 56.6 | -- |

**Table S5:** P-values of interaction term in linear model for chamber experiment response variable

|  | response variable |  |  |
| --- | --- | --- | --- |
|  | seedling height | total leaf length | relative height increment |
| maternal line | 0.89 | 0.58 | 0.67 |
| sampling site | 0.63 | 0.51 | 0.53 |

**Table S6:**
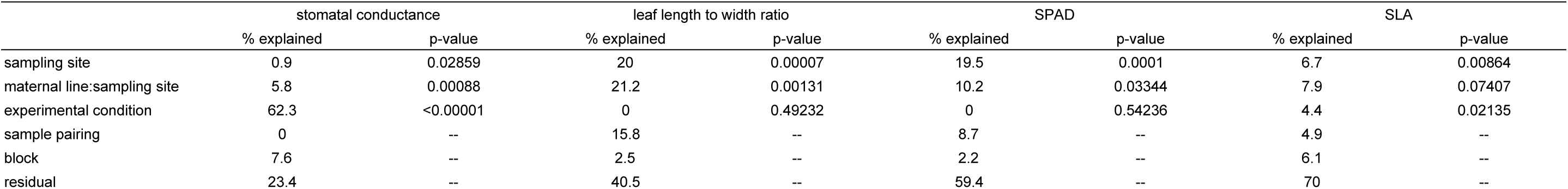
Percent of variation explained and p-values for non-interaction linear models for drought experiment

**Table S7:** P-values of interaction term in linear model for drought experiment

|  | response variable |  |  |  |
| --- | --- | --- | --- | --- |
|  | stomatal conductance | leaf length to width ratio | SPAD | SLA |
| maternal line | 0.13 | 0.47 | 0.56 | 0.27 |
| sampling site | 0.51 | 0.78 | 0.31 | 0.82 |

